# Comparing serial and parallel language input: Shared neural representations with distinct encoding strengths

**DOI:** 10.64898/2026.09.23.753812

**Authors:** Simone M. Krogh, Liina Pylkkänen

## Abstract

Language can be experienced in highly serial form, as in speech, or in fully parallel form, as in text, yet how our brains adapt to differences in seriality is not understood. Here, we held modality constant (visual) and examined whether processing mechanisms underlying grammaticality detection are similar under Rapid Serial Visual Presentation (RSVP) and Rapid Parallel Visual Presentation (RPVP). To identify shared neural representations of grammaticality, we trained a classifier on magnetoencephalography (MEG) responses to grammatical and ungrammatical three-word phrases in one presentation mode and tested its generalization to the other. Behaviorally, accuracy was higher and reaction times faster for grammatical vs. ungrammatical phrases across presentation modes. Neurally, cross-presentation decoding revealed shared representations of grammaticality at word 2 across train-test directions—albeit with different latencies and durations—and an early shared representation at word 3 when trained on RPVP. We take the bidirectional representations at word 2 to capture distinct but presentation-invariant combinatorial processes and the asymmetry at word 3 as reflecting stronger global structure encoding in RPVP.

## INTRODUCTION

Which aspects of language comprehension are shared across different degrees of seriality in the input? Serial presentation has enjoyed a privileged position in the study of the neurobiology of comprehension; speech is inherently serial, and decades of reading research has enforced seriality in the visual domain using Rapid Serial Visual Presentation (RSVP), which delivers words one-by-one. Thus our current understanding of the mechanisms underlying language comprehension has been shaped predominantly by studies using serial presentation (Friederici, 2002, 2011; Hickok & Poeppel, 2007; Pylkkänen, 2019).

In recent years, however, Rapid Parallel Visual Presentation (RPVP) has gained traction as an alternative way to present visual stimuli (Snell & Grainger, 2017). Here, short multi-word expressions are presented in their entirety for a few hundred milliseconds, with the absence of stimulus-internal temporal structure allowing linguistic computations to unfold in whatever order is natural for the system (Fallon & Pylkkänen, 2024). Instances of the so-called Sentence Superiority Effect show that the language system does indeed operate on the stimuli despite the rapid presentation: Grammatical sentences are identified faster and/or more accurately than word lists or scrambled sentences across a host of behavioral tasks (e.g., Snell & Grainger, 2017; Mirault et al., 2018; Pegado et al., 2021; Fallon & Pylkkänen, 2024). M/EEG studies have provided neural characterizations of the Sentence Superiority Effect, with activity diverging in left fronto-temporal areas as early as 130 ms post-stimulus onset and lasting several hundred milliseconds (Wen et al., 2019, 2021; Pegado et al., 2021; Dufau et al., 2024; Fallon & Pylkkänen, 2024; Flower & Pylkkänen, 2024, 2026a, 2026b; Krogh & Pylkkänen, 2025). Most recently, a similar effect was observed with phrases (Li & Pylkkänen, 2026), suggesting it to reflect general *structure* superiority.

RSVP and RPVP both have their advantages and disadvantages. The word-byword presentation of RSVP enables strict control of word processing, facilitating the detailed analysis of specific phenomena. To give just one example, a host of MEG studies has led to a detailed characterization of left anterior temporal lobe (LATL) activity increases at 200 ms associated with basic conceptual composition (e.g., Bemis & Pylkkänen, 2011, 2013; Zhang & Pylkkänen, 2015; Li & Pylkkänen, 2020; Parrish & Pylkkänen, 2022). However, in experimental settings, RSVP reading can be quite unnatural; despite the “rapid” in its name, it is typically presented at around 600 ms per word, far slower than natural silent reading with eye movements (∼230–250 ms per word; Brysbaert, 2019). Even within the RSVP literature, presentation rates as fast as 83 ms per word have been shown to allow comprehension and recall (Potter et al., 2014). As a result, slowly unfolding RSVP stimuli may elicit computations distinct from more natural language comprehension. For example, prediction, a central component of language processing (e.g., Lau et al., 2006, 2013; Brothers et al., 2015, 2023; Szewczyk & Schriefers, 2018; Yano, 2018; Kuperberg et al., 2020; Zhang et al., 2023), may be unnaturally enhanced given the large amount of time available to anticipate the next word. In contrast, RPVP eliminates prediction from temporally preceding context, allowing for clean tests of the topdown effects of grammatical knowledge. Results so far have revealed an early processing stage in which basic phrase structure influences neural signals, but not yet agreement, semantic roles, or verb argument structure (Fallon & Pylkkänen, 2024; Flower & Pylkkänen, 2024, 2026a; Dunagan et al., 2025; Krogh & Pylkkänen, 2025; Li & Pylkkänen, 2026).

Given the complementary advantages of serial and parallel language presentation, and the need to understand how the language system flexibly accommodates both, a key question is the extent to which stimuli with different temporal structures engage shared processing mechanisms. Here, we investigate the similarity of neural representations elicited by RSVP and RPVP using cross-temporal, cross-presentation decoding of MEG activity (King & Dehaene, 2014) in response to three-word-phrases (semantically designed to address questions about animacy, not reported here). Specifically, we trained classifiers to discriminate between neural responses to grammatical (e.g., *Arabian valley tiger*) and ungrammatical phrases (e.g., *valley Arabian tiger*) presented in one presentation mode, and then assessed classifier performance when the same phrases were presented via the other presentation mode. The main advantage of our cross-presentation temporal generalization approach is that it captures shared patterns: Successful classification requires high rates of similarity across representations found in training and test data and thus likely reflects similar computations induced by both presentation modes. Moreover, the use of temporal generalization, rather than standard time-resolved decoding, allows us to probe for temporally asynchronous effects. Neural activation profiles between stimuli presented with RSVP and RPVP are expected to temporally diverge given the inherent differences in stimulus delivery timing and input structure, and it is therefore possible that similar representations emerge with different latencies across presentation modes. In addition to cross-presentation decoding, we also report within-presentation decoding for RPVP and for each word position in the RSVP sequence. These analyses provide a reference against which to interpret cross-presentation generalization and characterize the temporal dynamics of grammaticality representations within each presentation mode.

Our study entertains a complex hypothesis space for the cross-presentation results, reflecting potential interactions between train-test directionality and serial word position (see Figure 1). If neural representations of grammatical and ungrammatical phrases are identical across presentation modes, we expect successful cross-presentation decoding regardless of train-test direction (H1). Alternatively, the same underlying representation may be encoded more strongly in one presentation mode than the other, leading to successful cross-presentation decoding when training on one presentation mode and testing on the other, but not vice versa. This could be the case for training on RPVP phrases and testing on RSVP phrases (H2) as well as for training on RSVP phrases and testing on RPVP phrases (H3). Since the three words in an RSVP phrase must be compared to the entire RPVP phrase one at a time, distinct patterns may emerge across word positions, consistent with different compositional subroutines at individual words. We expect shared representations of grammaticality to be particularly prominent at the second word, due to its central visual field position in RPVP, allowing us to probe cross-presentation representations at the exact point of composition (or the failure thereof).

**Figure 1:**
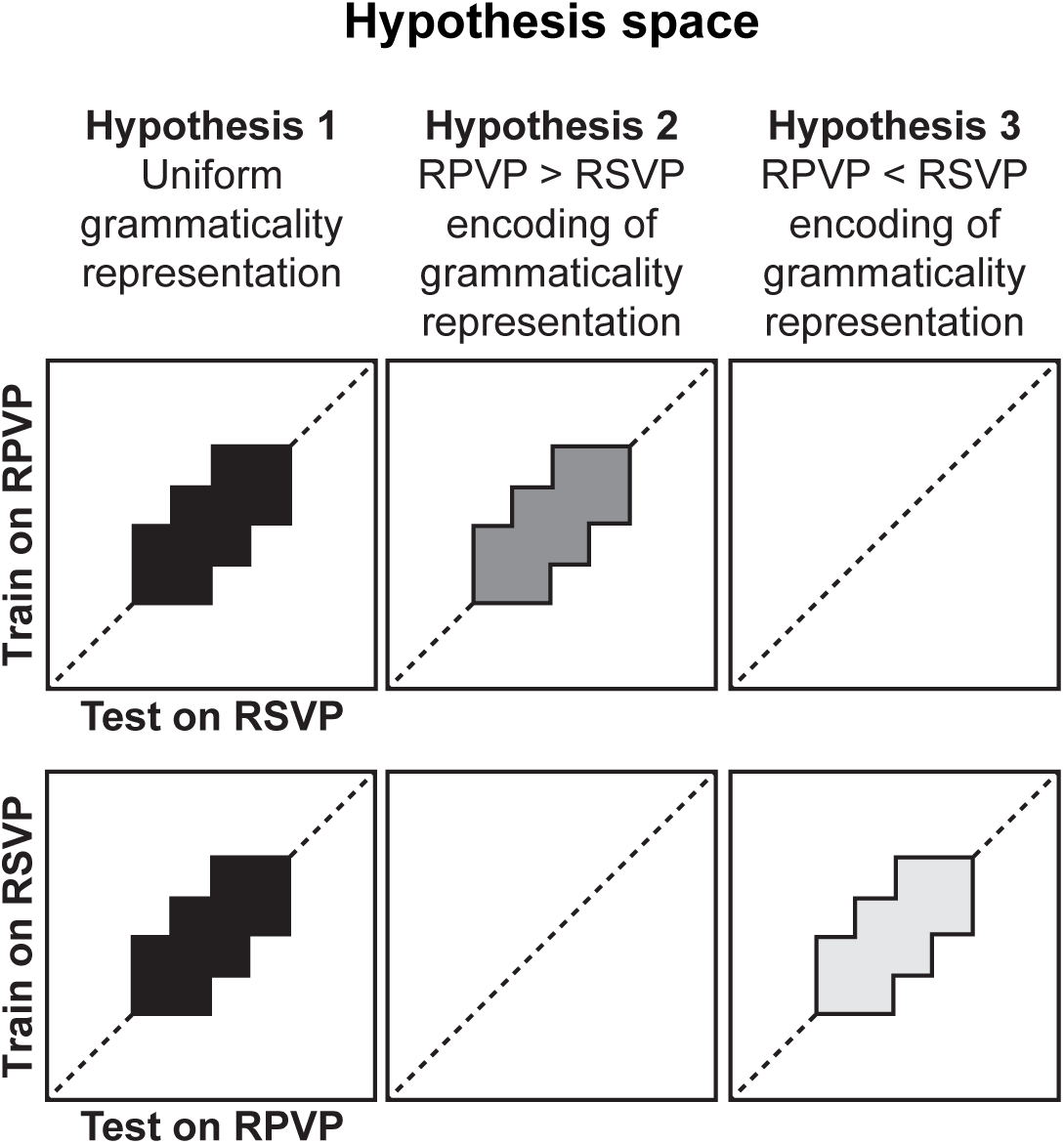
Hypotheses and their corresponding predictions for the cross-presentation decoding analyses. Neural representations of grammatical and ungrammatical phrases may be identical across presentation modes, resulting in successful decoding in either train-test direction (**H1**). Alternatively, sensitivity to grammaticality may be encoded more strongly in either RPVP (**H2**) or RSVP (**H3**). The different hypotheses need not apply uniformly to all word positions for RSVP. Diagonal clusters reflect shared, concurrent representations across presentation modes (illustrated here); off-diagonal clusters reflect shared but temporally shifted representations. RPVP: Rapid Parallel Visual Presentation; RSVP: Rapid Serial Visual Presentation.

## METHODS

### Participants

37 native speakers of English completed the experimental protocol. All participants were neurologically intact with normal or corrected-to-normal vision. Participants provided their informed written consent and were paid $15/hr for their participation. The study was approved by the Institutional Review Board (IRB) ethics committee of [redacted] (approval no.[redacted]). Eight participants were excluded from analyses (one for reporting a panic attack; one for sleepiness; six for excessive movement). This left 29 participants’ data in the final dataset (16 female, 9 male, 4 nonbinary; 20-40 years old, mean age ± SD: 26.2 years ± 5.1 years).

### Stimuli

Data reported in the present paper were obtained as part of a larger study that, in addition to characterizing phrase processing as a function of stimulus delivery technique, investigated the decodability of animacy in composed concepts. The study therefore employed a 2 × 2 × 2 factorial design with factors grammaticality (grammatical, ungrammatical), initial modifier type (geographical, material), and noun-noun order (location-animal, animal-location; see Table 1).

**Table 1:** Design table. The study employed a 2 × 2 × 2 factorial design (50 trials per condition), but grammaticality was the sole focus of this study. All stimuli were presented once using RSVP and once using RPVP.

| **Grammaticality** | **Initial modifier** | **Noun-noun order** | **Example** |
| --- | --- | --- | --- |
| grammatical | geographical | location-animal | ARABIAN VALLEY TIGER |
| grammatical | geographical | animal-location | ARABIAN TIGER VALLEY |
| grammatical | material | location-animal | PLASTIC VALLEY TIGER |
| grammatical | material | animal-location | PLASTIC TIGER VALLEY |
| ungrammatical | geographical | location-animal | VALLEY ARABIAN TIGER |
| ungrammatical | geographical | animal-location | TIGER ARABIAN VALLEY |
| ungrammatical | material | location-animal | VALLEY PLASTIC TIGER |
| ungrammatical | material | animal-location | TIGER PLASTIC VALLEY |

For the noun-noun order manipulation, 25 animals and 25 locations matched for length and other lexical characteristics were identified and paired in two plausible but novel combinations (e.g., *tiger* was paired with *valley* and *desert*) as verified by transition probabilities of 0% in COCA (Davies, 2008) and < 0.07% in iWeb (Davies, 2018) for both possible word orders (*valley tiger* or *tiger valley*). For the manipulation of initial modifier type, each of the 100 pairs was modified by one of ten geographical placenames (e.g., *Arabian*) and one of ten length-matched constitutive materials (e.g., *plastic*), resulting in 200 grammatical three-word phrases. For the manipulation of grammaticality, ungrammatical phrases were formed by swapping the first two words in grammatical phrases (grammatical: *Arabian valley tiger*; ungrammatical: *valley Arabian tiger*) for a total of 400 trials with 50 trials in each of the eight conditions. Crucially, this made grammatical and ungrammatical phrases distinguishable at the very first word. All stimuli were presented in all caps to avoid confounding effects from geographic placenames conventionally being capitalized. A full three-word phrase subtended a visual angle between 5.85° and 8.78° (mean ± SD: 7.29°± 0.65°) whereas individual words subtended a visual angle between 1.46° and 3.29° (mean ± SD: 2.15°± 0.50°). Results obtained from the manipulation of initial modifier type and noun-noun order will be discussed elsewhere.

### Experimental design

The experimental protocol included four components: phrases presented in RSVP (4 blocks), phrases presented in RPVP (4 blocks), and single-word trials used to address animacy decodability questions (10 obligatory blocks and 10 optional blocks; results are discussed in separate reports). The 18 obligatory blocks were presented in a semi-structured, pseudorandomized manner following which participants could choose to do the 10 optional blocks. Here, we focus only on the two phrasal components in which participants read three-word phrases word-by-word (RSVP) or all-at-once (RPVP). Presentation mode alternated across blocks, with the order of first presentation mode counterbalanced across participants. Since all 400 three-word phrases were presented in both RSVP and RPVP, presentation order was counterbalanced such that trials appearing early in one presentation mode appeared late in the other.

Regardless of presentation mode, a trial always began with a fixation cross (200 ms) and a blank screen (200 ms; Figure 2). In RSVP, each of the three words in the phrase (target stimulus) was then presented for 300 ms followed by a 500 ms blank screen. In RPVP, the entire three-word phrase (target stimulus) was shown for 300 ms and followed by a 500 ms blank screen. Following prior RPVP-MEG studies, we used a simple matching task that has previously been shown to elicit Structure Superiority Effects, providing evidence for an immediate and automatic effect of top-down grammatical knowledge despite the shallow nature of the task (Fallon & Pylkkänen, 2024; Flower & Pylkkänen, 2024, 2026a, 2026b; Dunagan et al., 2025; Krogh & Pylkkänen, 2025; Li & Pylkkänen, 2026). Regardless of presentation mode, trials thus concluded with a three-word task stimulus shown for 300 ms followed by a blank screen until button press. The task stimulus was either identical to the target stimulus (a match trial) or differed by one word (a mismatch trial). The position of the swapped word in the task stimulus was randomized across trials but the type of word (animal, location, geographical modifier, material modifier) and length were kept constant between target and task stimuli.

**Figure 2:**
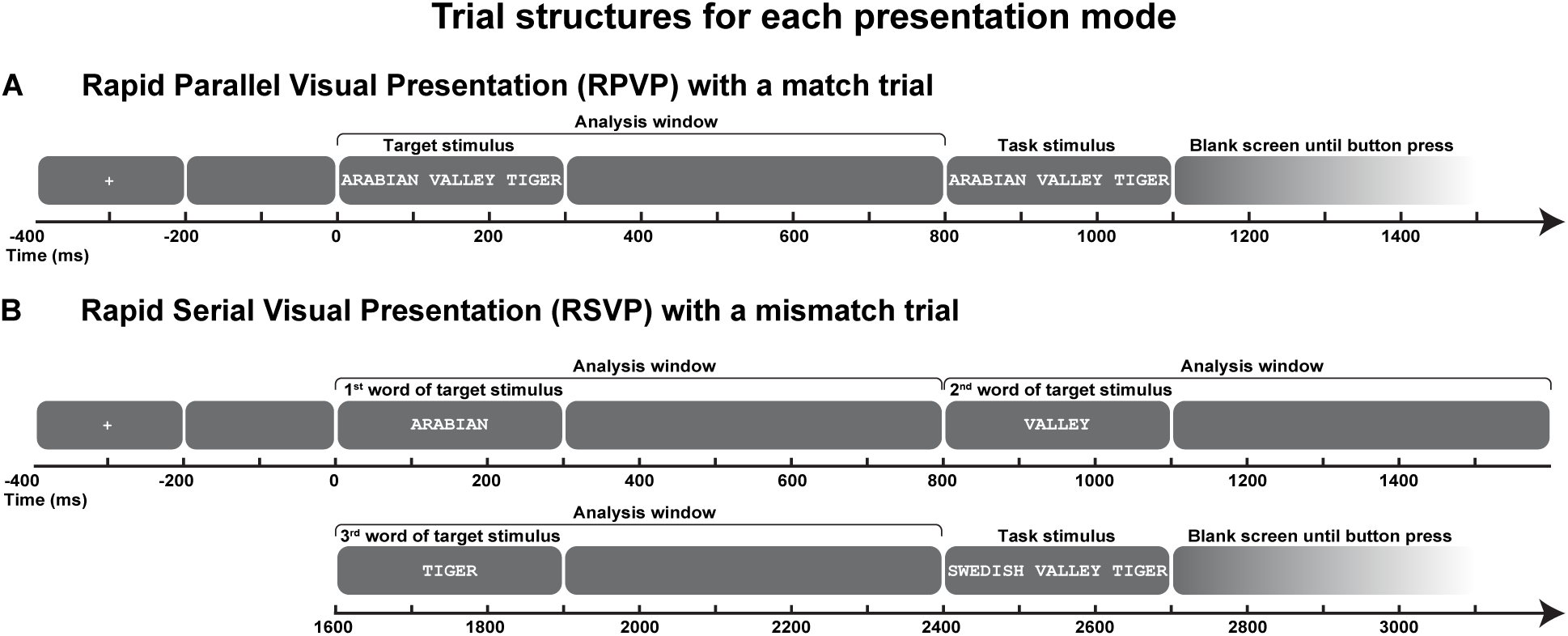
Trial structures for RPVP and RSVP. In RPVP (**A**), a full three-word phrase was presented for 300 ms followed by a blank screen for 500 ms. In RSVP (**B**), each word was presented in a 300 ms on, 500 ms off-sequence. Each trial was followed by a three-word target stimulus that was either identical to (match trial) or differed by one word from (mismatch trial) the target stimulus. Here, the RPVP trial structure illustrates a match trial and the RSVP trial structure a mismatch trial. The data submitted to analysis only included the target stimulus (the entire three-word phrase in RPVP and individual words in RSVP), not the task stimulus. RPVP: Rapid Parallel Visual Presentation; RSVP: Rapid Serial Visual Presentation.

### Procedure

Upon arrival to the lab, participants gave informed consent and filled out a brief demographic questionnaire. Then, 3D digitizations of participants’ head shapes were recorded with a Polhemus FastSCAN laser scanner (Polhemus VT, USA) to enable coregistration of the MEG data with the FreeSurfer *fsaverage* brain (Fischl, 2012). The digitizations included five future marker coil placements and the location of three fiducial landmarks (nasion, left and right tragi). After a brief instruction period and practice sessions for all experimental components, participants were set up in the magnetically shielded room hosting the MEG machine and completed the experiment in supine position.

### MEG data acquisition and preprocessing

Continuous MEG data were recorded using a whole-head, 157-channel axial gradiometer system (Kanazawa Institute of Technology, Kanazawa, Japan) at a sampling rate of 1000 Hz with an online bandpass filter of 0.1–200 Hz. Participants’ head positions relative to the MEG sensors were recorded before and after the experiment by means of marker coils. To ensure timing precision, a photodiode was used to measure stimulus-to-trigger delays.

In an initial preprocessing step, the Continuously Adjusted Least-Squares Method (Adachi et al., 2001) as implemented in the MEG160 software (Yokogawa Electrical Corporation and Eagle Technology Corporation, Japan) was used to noise-reduce the MEG data based on three reference channels. Subsequent data preprocessing was undertaken in the Python computing environment using the *MNE Python* v1.10 package (Gramfort et al., 2014) and consisted of the following steps. After applying a 1 to 40 Hz offline bandpass filter (with the high-pass filter being necessary to attenuate NYC environmental noise), we removed and interpolated two known broken channels as well as recording-specific saturated channels (mean: 1.45 additional bad channels) based on data from neighboring channels. Then, Independent Component Analysis (ICA) was used to separate neural data from ocular, cardiac, and environmental artifacts. Epochs comprising the target stimulus and the ensuing blank screen were extracted for the three-word phrases presented in RPVP (800 ms) and in RSVP (2400 ms) and baseline-corrected using the 100 ms preceding the onset of (the first word in) the target stimulus. Individual epochs were rejected based on a 3000fT peak-to-peak amplitude threshold, technical issues flagged during acquisition, or behavioral response anomalies (e.g., incorrect responses or response times more than 3 SDs from participant means). Finally, RSVP epochs were split into three 800 ms epochs (RSVP word 1, RSVP word 2, RSVP word 3) to allow for direct comparison with RPVP epochs. See Figure 3 for grand averages of the left-hemisphere sensors in RPVP and at each word in RSVP. Note that grand averages for RSVP words 2 and 3 are plotted with the 100 ms baseline of RSVP word 1 to facilitate direct visual comparison, though baselines were never included in the statistical analysis.

**Figure 3:**
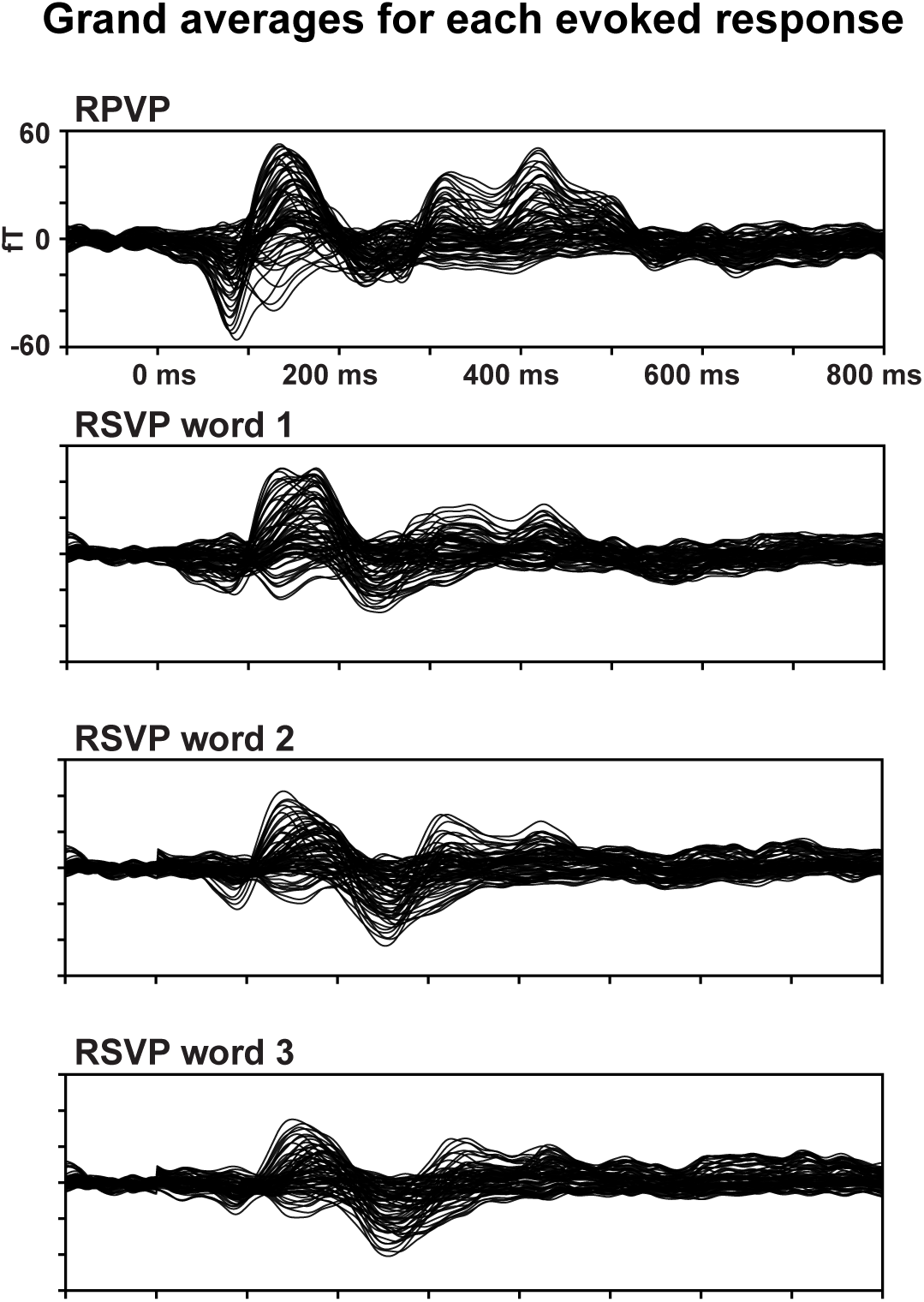
Grand averages of left hemisphere sensors across all participants for RPVP phrases and each word in RSVP. Despite some amplitude differences, overall waveforms were remarkably similar across RPVP and each of the RSVP words. The baseline of RSVP word 1 is repeated for RSVP words 2 and 3 for visual comparison; baselines were not included in any statistical analyses. RPVP: Rapid Parallel Visual Presentation; RSVP: Rapid Serial Visual Presentation.

For our spatial decoding analyses, we additionally produced single-trial source time courses. This involved coregistering the FreeSurfer *fsaverage* template brain to each participant’s digitized head shape and fiducials, which was then transformed to a source-space mesh of 2,562 vertices per hemisphere (ico-4 spacing). Forward solutions were computed via the Boundary Element Model (BEM), and channel noise-covariance matrices were derived from the 100 ms pre-stimulus interval across all trials. Forward solutions and channel noise-covariance matrices were used to calculate subject-specific inverse solutions, applying an SNR of 3. The entire procedure yielded L2 minimum norm source estimates, which were subsequently noise-normalized to generate Dynamic Statistical Parameter Maps (dSPMs; Dale et al., 2000).

### Statistical analysis

#### Behavioral data analysis

Our behavioral analyses focused on detecting sensitivity to grammaticality within and across presentation modes. The data were first cleaned of responses with reaction times more than 3 SDs from a participant’s overall mean and then subjected to (generalized) linear mixed effects regression models with grammaticality (grammatical, ungrammatical), presentation mode (RPVP, RSVP), and response type (match trial, mismatch trial) as fixed effects alongside all possible interaction terms and the maximally converging random effects structures. Response type was included as a fixed effect since match and mismatch trials were expected to differ in processing difficulty, a prediction that has been borne out in prior research using the matching task with RPVP (Fallon & Pylkkänen, 2024; Flower & Pylkkänen, 2024, 2026a; Krogh & Pylkkänen, 2025; Li & Pylkkänen, 2026). The linear mixed-effects regression model of the reaction time data took log-transformed reaction time data (incorrect responses excluded) as the dependent variable and had by-participant and by-item intercepts. The generalized linear mixed-effects regression model of the accuracy data (incorrect responses included) took accuracy as the dependent variable and had by-participant and by-item intercepts as well as uncorrelated by-participant random slopes for response type, presentation mode, and their interaction. All *p* values were generated via likelihood ratio tests, and pairwise comparisons were corrected using a Tukey adjustment. Analyses were conducted using the *lme4* (Bates et al., 2015) and *afex* (Singmann et al., 2023) packages in R (v4.5.2) and RStudio (v2026.01).

Words in different positions (left, middle, or right in RPVP; early, mid, or late in RSVP) may receive differential attention due to differences in perceptual prominence or working memory load. To evaluate this possibility, we ran two control analyses restricted to mismatch trials, with the position of the swapped word (first, second, third) as a fixed effect alongside grammaticality and presentation mode. If word positions were differentially processed, this should be reflected on the mismatch trials. The reaction time model again only had by-participant and by-item intercepts while the accuracy model also had an uncorrelated by-participant random slope for presentation mode.

#### Generalization across time and presentation modes

For our primary decoding analyses, we trained unique time-resolved decoders to discriminate between neural responses corresponding to grammatical and ungrammatical phrases in one presentation mode and tested their generalization to the other. This was done for each train-test combination of the RPVP epochs and the three RSVP epochs, resulting in a total of six distinct analyses.

To enable faster computing and increase the signal-to-noise ratio (Grootswagers et al., 2017), we (1) downsampled epochs by averaging non-overlapping bins of 5 ms, (2) averaged two randomly sampled epochs from the same experimental condition without replacement, and (3) reduced the sensor space with Principal Component Analysis (PCA), transforming the 75 left-hemisphere sensors (midline sensors excluded) into 40 principal components explaining at least 96.78% of the variance. We restricted our analyses to the left hemisphere based on the left-lateralized results of earlier RPVP-MEG studies (Dufau et al., 2024; Fallon & Pylkkänen, 2024; Flower & Pylkkänen, 2024, 2026a, 2026b; Krogh & Pylkkänen, 2025; Li & Pylkkänen, 2026).

For each participant, the following pipeline was performed to produce temporal generalization matrices. The 40 principal components, scaled to unit variance, were fed into a logistic regression classifier with l2 regularization. Regularization strength (C) was optimized via a grid search on a logarithmic scale between 1e^−4^ and 1e^4^ using stratified 5-fold cross-validation on the training data. Mechanistically, classification was done by training and testing separate unique classifiers on pairs of time points using 100-ms windows (with edge padding), stepped at 10-ms intervals throughout the 0–800 ms epoch. Notably, we employed a 5-fold cross validation on our cross-presentation data sets, emulating that used for within-presentation decoding (e.g., training and testing on RPVP phrases) to facilitate comparisons. In practice, classifiers were thus trained on 80% of the training set (e.g., RPVP) and tested on 20% of the test set (e.g., RSVP word 1). This procedure was repeated across five folds, with the resulting scores averaged. Since data preprocessing often left different trial counts for RPVP and RSVP trials, we randomly subsampled the larger set to ensure that grammatical and ungrammatical trial counts were matched separately across presentation modes.

#### Generalization across time but within presentation mode

As a control analysis against which to interpret cross-presentation results, we performed within-presentation decoding for RPVP and each word position in the RSVP sequence. This yielded ten additional analyses: one for the RPVP data, and nine representing all possible train-test combinations across the three RSVP word positions. Implementation details are otherwise as described in *Generalization across time and presentation modes.* While the training and test sets for these within-presentation analyses already had identical trial counts, we subsampled them to match the trial counts used in the cross-presentation analyses. Doing so controlled the dataset size and thus ensured an appropriately powered comparison between the within- and cross-presentation analyses.

#### Group-level statistical testing of temporal generalization analyses

Above-chance decoding at the group-level was assessed using cluster-based permutation tests (Maris & Oostenveld, 2007). For each analysis, group-level decoding accuracy at each pair of time points was evaluated against chance (0.5) using a one-tailed, one-sample *t* test, with resulting *t* values thresholded at those corresponding to an uncorrected *p* value of 0.05. Temporally adjacent pairs of time points were clustered using summed *t* values, and cluster significance was determined with Monte Carlo simulations (10,000 permutations). Only clusters with a size of ≥ 2 points (equal to a minimum cluster duration of 20 ms) and a cluster *p* value ≤ 0.05 were retained.

We performed two group-level analyses: one in a targeted 100–400 ms and one in the full 100–700 ms viable window (accounting for sliding window edge padding). The 100–400 ms window allowed us to isolate a time period previously associated with combinatory responses across both MEG-RSVP (for a review, see Pylkkänen, 2019) and MEG-RPVP (Dufau et al., 2024; Fallon & Pylkkänen, 2024; Flower & Pylkkänen, 2024, 2026b; Krogh & Pylkkänen, 2025; Li & Pylkkänen, 2026) studies. However, because focusing exclusively on this restricted interval limits the detection of temporally asynchronous effects, we also probed the broader 100–700 ms window. Note that expanding the temporal dimensions scales the 2D analysis matrix quadratically, which drastically raises the cluster-size threshold required for significance, and effects in this larger window need to be considerably more robust to survive the multiple-comparisons correction of the cluster permutation test. The temporal extent of any significant cluster should be interpreted with some caution since clusters were formed using uncorrected *t* values (Sassenhagen & Draschkow, 2019). MEG analysis scripts were written in Cursor, an AI-assisted code editor used for limited code writing, editing, and debugging with both automatic model routing (Auto) and manually selected Claude Sonnet and Opus models (4.5 and 4.6). All aspects of the decoding analyses were performed in Python using *MNE-Python* (v1.0.3) and *scikit-learn* v1.6.1 (Pedregosa et al., 2011).

#### Spatial decoding analyses across and within presentation modes

To identify the principal neural generators of the effects observed in our sensor-space temporal decoding analyses, we followed up with spatial decoding analyses at the source level (for similar approaches, see e.g., Gwilliams & King (2020) and Zuanazzi et al. (2024)). For each significant cluster identified in the temporal decoding analyses, we performed 10mm-radius searchlight analyses at each neural source, using the temporal neural patterns within 50 ms × 50 ms train-test tiles from the temporal generalization matrix. Tiles were uniformly placed on the temporal clusters with the first tile being 101–150 ms × 101–150 ms. Such grid-based partitioning guaranteed a standardized framework for comparing heterogeneous clusters observed across the temporal decoding analyses but was at the expense of maximal coverage of individual temporal clusters. We excluded temporal tiles with less than 20% coverage of significant clusters to ensure our spatial analysis focused on regions with substantial temporal overlap. The pipeline otherwise mirrored that described in *Generalization across time and presentation modes*.

#### Group-level statistical testing of spatial decoding analyses

Consistent with our temporal generalization analyses, we employed cluster-based permutation tests to assess group-level significance at each temporal tile (see *Group-level statistical testing of temporal generalization analyses*). We restricted our analyses to the language mask used in Fallon & Pylkkänen’s (2024) MEG-RPVP study. This included the left temporal and parietal lobes, the anterior portion of the occipital lobe (BA 19) as well as ventromedial and prefrontal areas (left BAs 10, 11, 44, 45, and 47). Because our spatial decoding analyses were guided by previously identified temporal clusters, the resulting *p* values are statistically non-independent and therefore only serve to characterize an effect’s spatial distribution rather than provide primary inferential evidence. Furthermore, it is important to note that temporal decoding analyses rely on fine-grained spatial patterns; as a result, individual sources within a cluster may contribute critically to pattern discriminability without necessarily exhibiting significant effects in isolation.

## RESULTS

### Behavioral sensitivity to grammaticality across presentation modes

We found clear processing advantages for grammatical over ungrammatical phrases, with main effects of grammaticality for both reaction time (*p* < 0.001) and accuracy (*p* < 0.001) as seen in Figure 4 and Table 2. In other words, participants responded faster and more accurately to grammatical phrases (815.65 ± 353.42 ms; 92.35 ± 6.46%) as compared to ungrammatical phrases (835.33 ± 359.45 ms; 89.98 ± 7.09%) across the entire dataset. In the case of reaction time, grammaticality interacted with presentation mode (*p* < 0.001), and while pairwise comparisons revealed the processing advantage for grammatical over ungrammatical phrases to be significant only for RSVP (*p* < 0.001), it remained marginally significant for RPVP (*p* = 0.0591). In the case of accuracy, grammaticality participated in a two-way interaction with response type as well as a complete three-way interaction. Decomposition of the latter revealed processing advantages for grammatical over ungrammatical phrases in all pairwise comparisons other than RSVP match trials (*p* = 0.9183; all other *p* < 0.0045).

**Figure 4:**
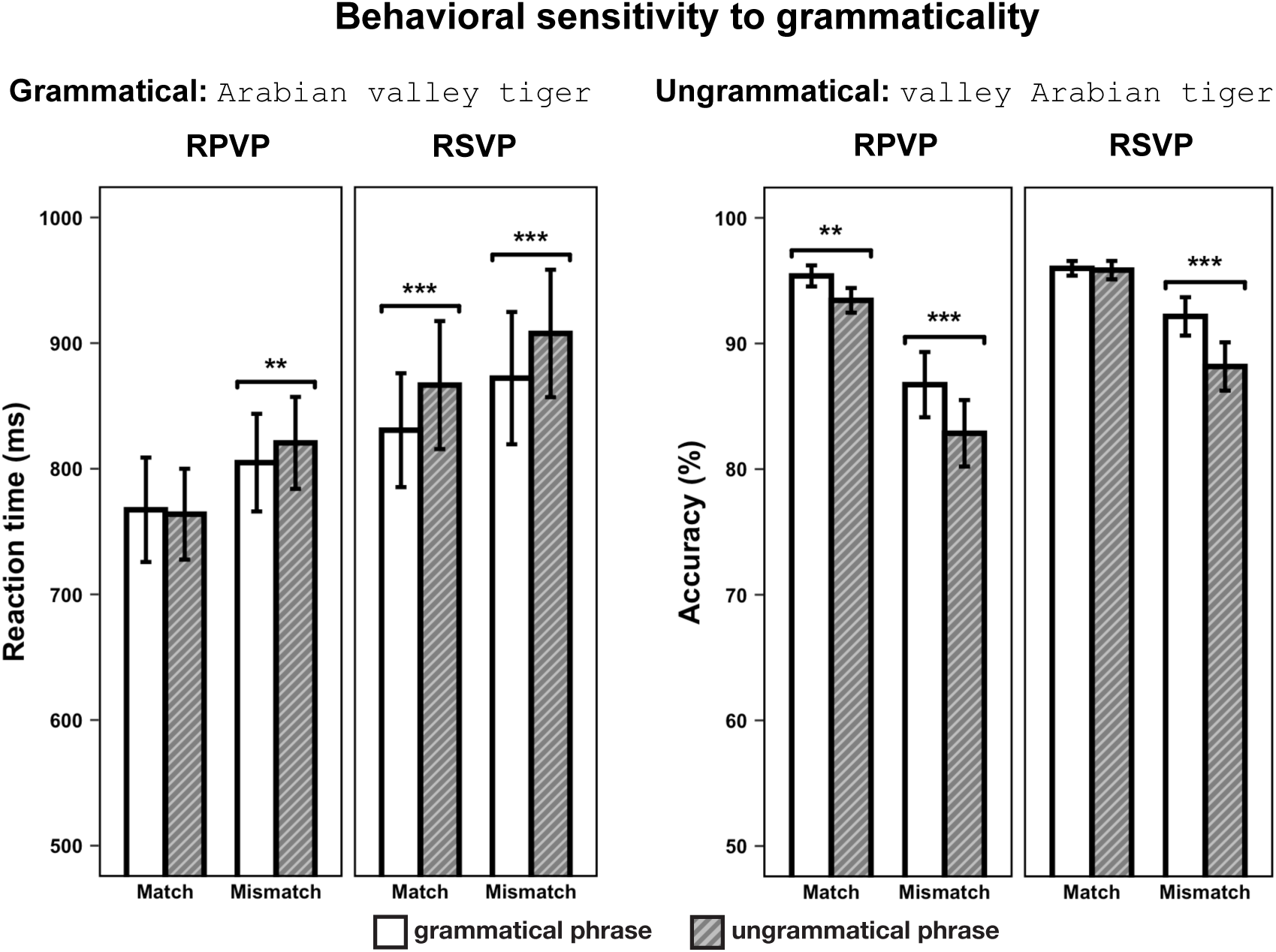
Behavioral sensitivity to grammaticality. Grammatical phrases elicited faster reaction times and higher accuracy than ungrammatical phrases, regardless of presentation mode. RPVP: Rapid Parallel Visual Presentation; RSVP: Rapid Serial Visual Presentation.

**Table 2:** Likelihood ratio tests of sensitivity to grammaticality. Log-transformed reaction times (incorrect responses excluded) were modeled using linear-mixed effects regression while accuracy was modeled using generalized linear mixed-effects regression.

| **Effect** | **df** | **Reaction time** |  | **Accuracy** |  |
| --- | --- | --- | --- | --- | --- |
|  |  | **$\chi^2$** | ***p* value** | **$\chi^2$** | ***p* value** |
| grammaticality | 1 | 23.37 | < 0.001 | 16.72 | < 0.001 |
| presentation mode | 1 | 527.49 | < 0.001 | 17.59 | < 0.001 |
| response type | 1 | 307.43 | < 0.001 | 21.70 | < 0.001 |
| grammaticality × presentation mode | 1 | 12.51 | < 0.001 | 0.88 | 0.349 |
| grammaticality × response type | 1 | 2.95 | 0.086 | 4.98 | 0.026 |
| presentation mode × response type | 1 | 4.19 | 0.041 | 1.69 | 0.193 |
| grammaticality × presentation mode × response type | 1 | 1.41 | 0.235 | 5.19 | 0.023 |

We also observed main and interaction effects of response type and presentation mode across both reaction time and accuracy (all *p* < 0.001). Unsurprisingly, participants responded faster and more accurately to match trials than mismatch trials. Interestingly, however, the effects of presentation mode on reaction times and accuracies differed; participants responded faster to phrases presented with RPVP but had higher accuracies when phrases were presented with RSVP.

Our control analyses examining the effect of the position of the swapped word on mismatch trials found a main effect of word position for reaction time (*p* = 0.026) but not for accuracy (*p* = 0.204). Post-hoc pairwise comparisons of the reaction time main effect revealed faster processing of word 2 mismatches relative to word 3 mismatches (*p* = 0.0254), with none of the other pairwise comparisons reaching significance (*p* > 0.122). Word position did not participate in any interactions. See Figure 5 and Table 3 for more details.

**Figure 5:**
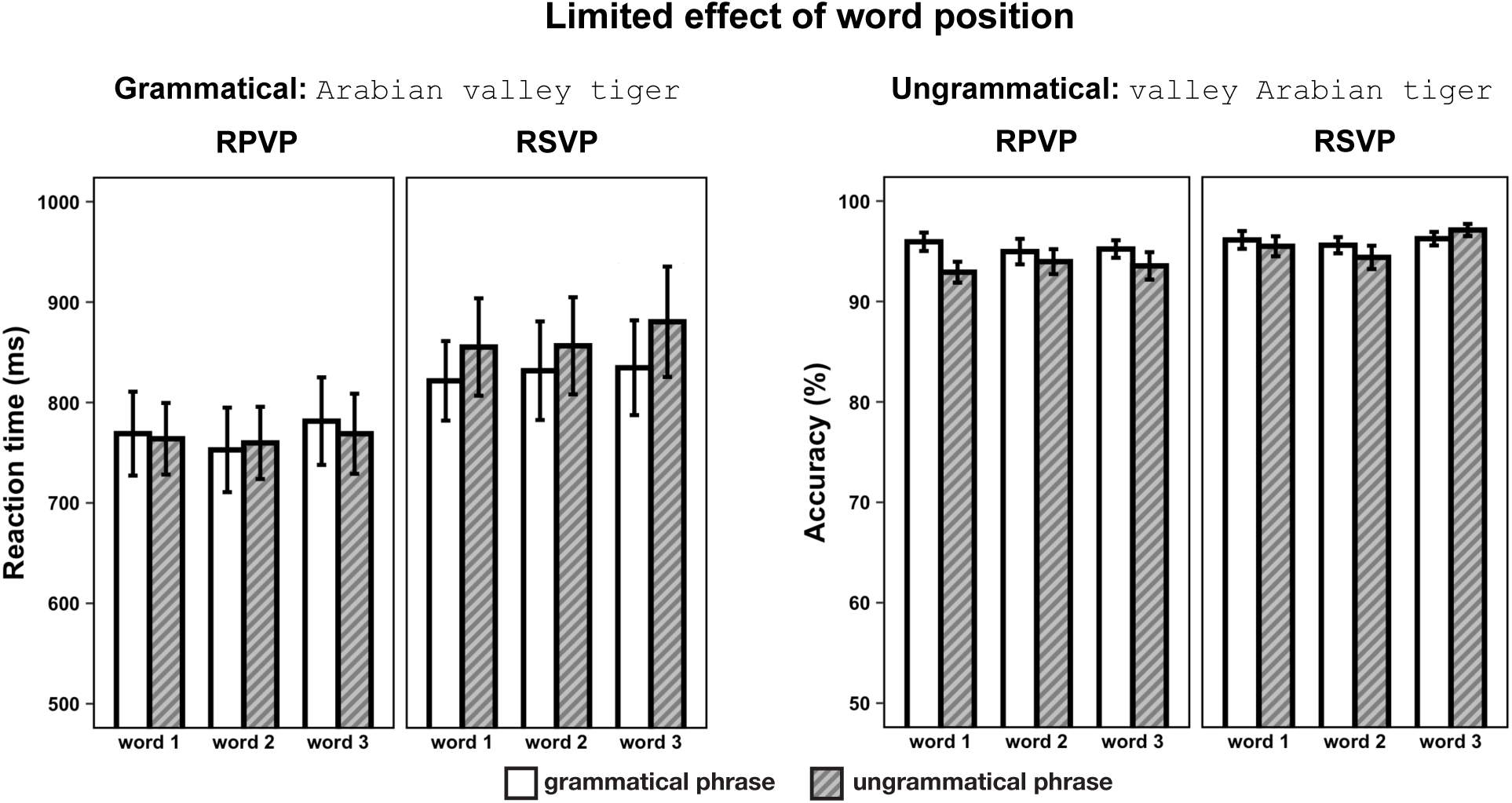
Limited effect of swapped word position in mismatch trials. Mismatch trials randomly swapped one of the three words between target and task stimuli (left, middle, or right in RPVP; early, mid, or late in RSVP), making it possible to investigate whether each word position was processed equally by participants. Except for word 2 having faster reaction times than word 3 across presentation modes, no evidence for differential processing was found. RPVP: Rapid Parallel Visual Presentation; RSVP: Rapid Serial Visual Presentation.

**Table 3:** Likelihood ratio tests investigating the impact of word position in mismatch trials. Log-transformed reaction times (incorrect responses excluded) were modeled using linear-mixed effects regression while accuracy was modeled using generalized linear mixed-effects regression.

| **Effect** | **df** | **Reaction time** |  | **Accuracy** |  |
| --- | --- | --- | --- | --- | --- |
|  |  | **$\chi^2$** | ***p* value** | **$\chi^2$** | ***p* value** |
| grammaticality | 1 | 7.73 | 0.005 | 4.08 | 0.043 |
| presentation mode | 1 | 282.63 | < 0.001 | 6.08 | 0.014 |
| word position | 2 | 7.30 | 0.026 | 3.18 | 0.204 |
| grammaticality × presentation mode | 1 | 9.26 | 0.002 | 3.28 | 0.070 |
| grammaticality × word position | 2 | 0.22 | 0.898 | 1.41 | 0.494 |
| presentation mode × word position | 2 | 1.05 | 0.590 | 2.78 | 0.250 |
| grammaticality × presentation mode × word position | 2 | 1.88 | 0.391 | 2.70 | 0.259 |

### Shared neural sensitivity to grammaticality across presentation modes, with distinct word position-profiles

Three of our six cross-presentation decoding analyses yielded significant clusters: bidirectional decodability at the second word as well as when training on RPVP and testing on RSVP word 3 (Figure 6). Specifically, training on RPVP and testing on RSVP word 2 produced a significant early cluster in the 100–400 ms window (train: ∼120–330 ms; test: ∼240–360 ms; *p* = 0.046) whereas the opposite train-test direction yielded a significant late cluster in the 100–700 ms window (train: ∼330–520 ms; test: ∼310–530 ms; *p* = 0.0382). Additionally, a significant early cluster arose from training on RPVP and testing on RSVP word 3 in the 100–400 ms window (train: ∼130–250 ms; test: ∼230–350 ms; *p* = 0.0472). Conversely, a marginal early cluster arose from training on RSVP word 3 and testing on RPVP in the 100–400 ms window (train: ∼310–400 ms; test: ∼100–230 ms; *p* = 0.0704). The remaining cross-presentation analyses were not significant (100–400 ms: all *p* ≥ 0.1; 100–700 ms: all *p* ≥ 0.16).

**Figure 6:**
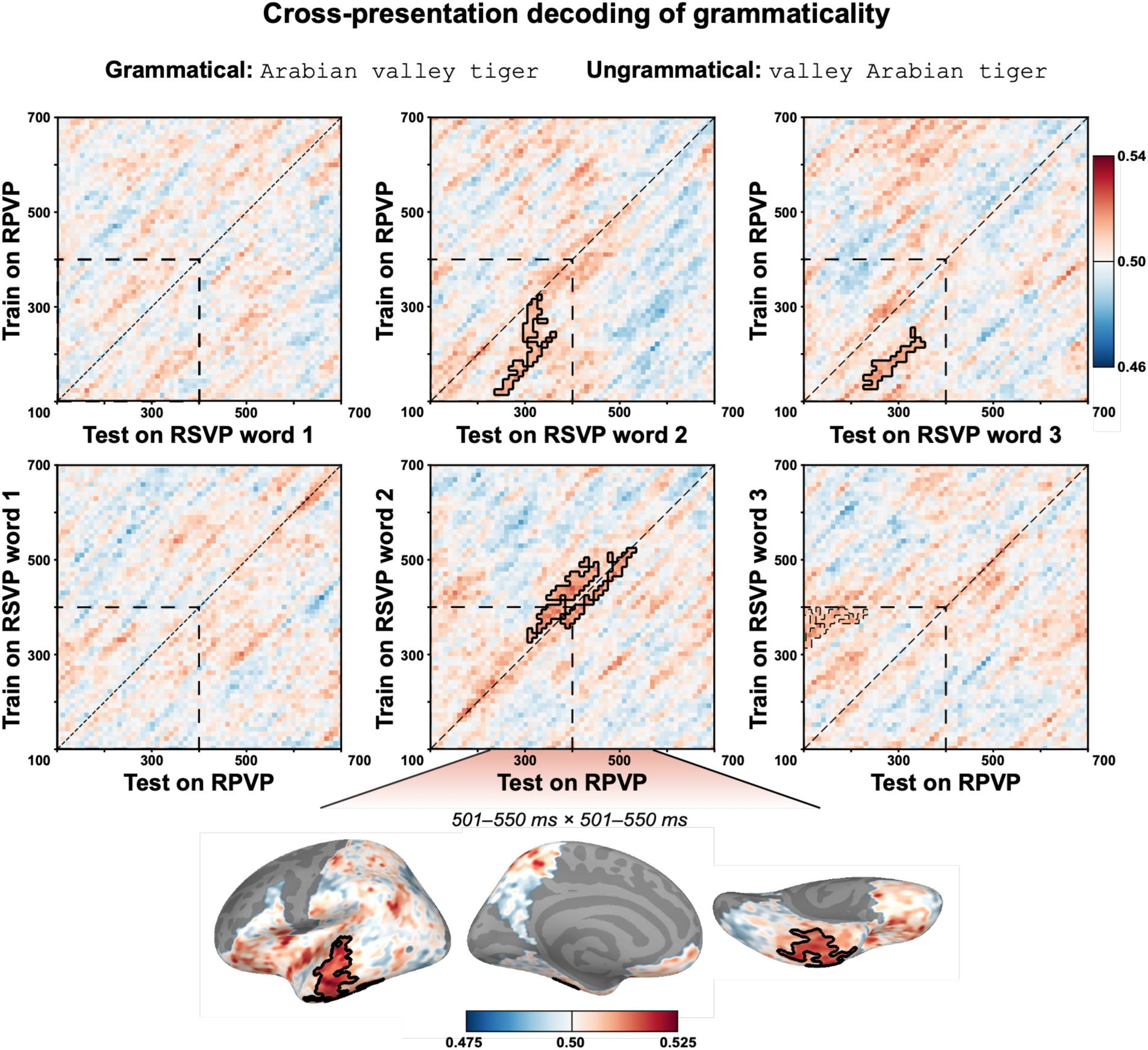
Cross-presentation temporal decoding of grammaticality. Classification was successful in both train-test directions at RSVP word 2, even though the clusters had different latencies and durations. Training on RSVP word 2 and testing on RPVP produced a spatial cluster with an anterior temporal localization. Additionally, an early cluster emerged when trained on RPVP and tested on RSVP word 3, with very similar timings as when training on RPVP and testing on RSVP word 2. Temporal train-test time windows were 100–400 ms and 100–700 ms. Solid cluster outline: *p* < 0.05; dashed cluster outline: 0.05 < *p* < 0.1. RPVP: Rapid Parallel Visual Presentation; RSVP: Rapid Serial Visual Presentation.

Follow-up spatial decoding analyses in the temporal clusters revealed one spatial cluster, namely when training on RSVP word 2 and testing on RPVP in the 501–550 ms × 501–550 ms (*p* = 0.0037) tile. This spatial cluster had an anterior temporal location, including both lateral and ventral areas. No spatial results arose from the temporal clusters associated with training on RPVP and testing on either RSVP word 2 or 3. A complete overview of cross-presentation clusters can be found in the Supplementary Materials (Table S1).

### Robust neural sensitivity to grammaticality within serial and parallel presentation modes

Several of our within-presentation decoding analyses, which served as control analyses against which to evaluate the cross-presentation results, yielded significant temporal clusters. Training and testing on the same type of linguistic input returned clusters with concurrent train-test times for RPVP, RSVP word 1, and RSVP word 2, but not RSVP word 3, although the latency and duration of these clusters varied substantially. As for generalization across RSVP word positions, we only observed successful generalization between RSVP words 2 and 3.

While spatial localization of such temporal clusters was not always possible, successful localizations typically followed a posterior-to-anterior progression: The earliest effects (starting at ∼100 ms) localized to occipital or posterior ventral cortex whereas most later effects (from ∼250 ms onwards) recruited temporoparietal, anterior temporal, and inferior frontal areas. More details on the ten within-presentation analyses are provided below. See Figure 7 for RPVP within-presentation decoding results, and Figure 8 for RSVP within-presentation decoding results. Complete overviews of temporal and spatial clusters are provided in the Supplementary Materials (Table S2 and S3).

**Figure 7:**
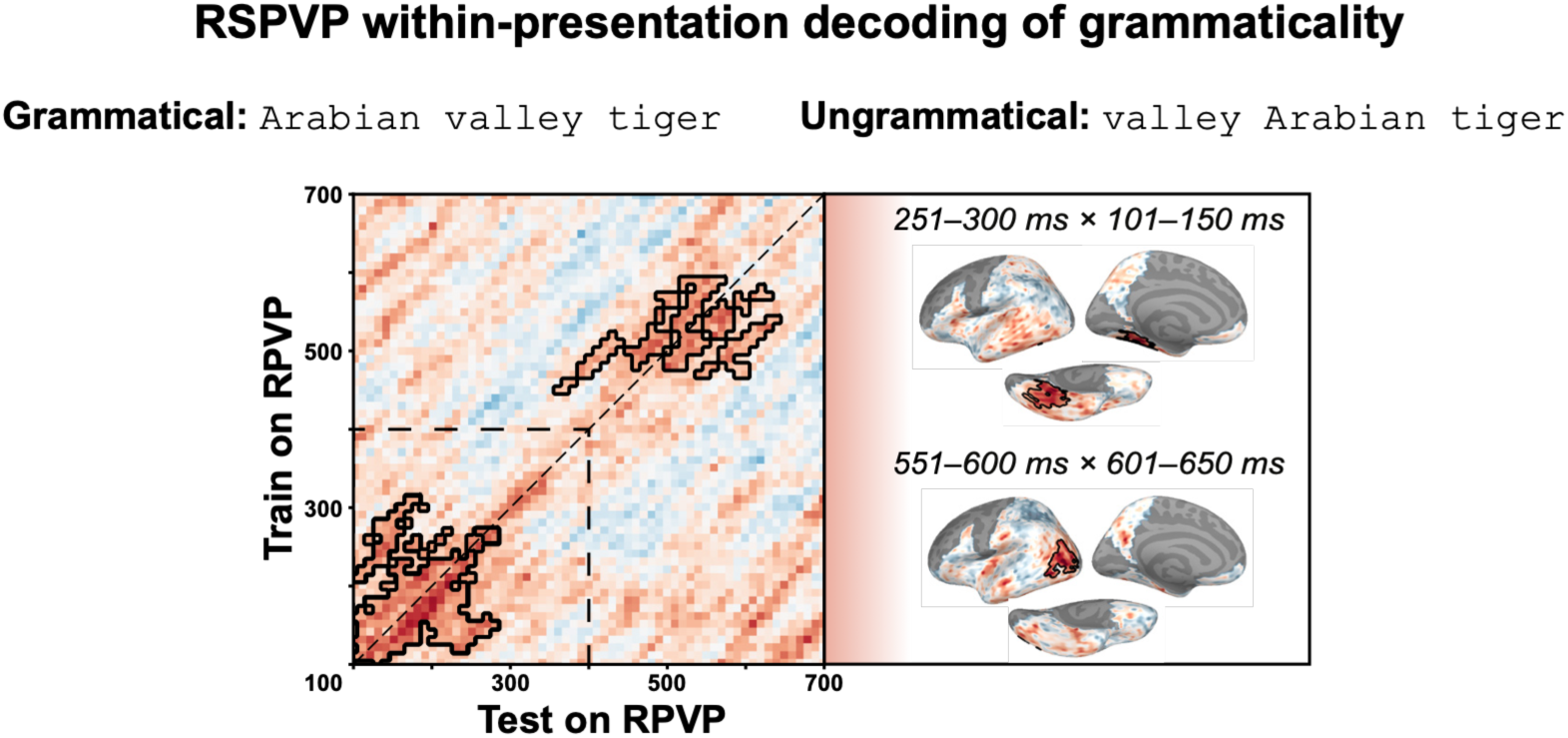
RPVP within-presentation decoding of grammaticality. Classifier training and testing on RPVP trials produced an early and a late temporal cluster, which recruited posterior-ventral temporal and occipital sources. Temporal train-test time windows were 100–400 ms and 100–700 ms. Solid cluster outline: *p* < 0.05; dashed cluster outline: 0.05 < *p* < 0.1. RPVP: Rapid Parallel Visual Presentation.

**Figure 8:**
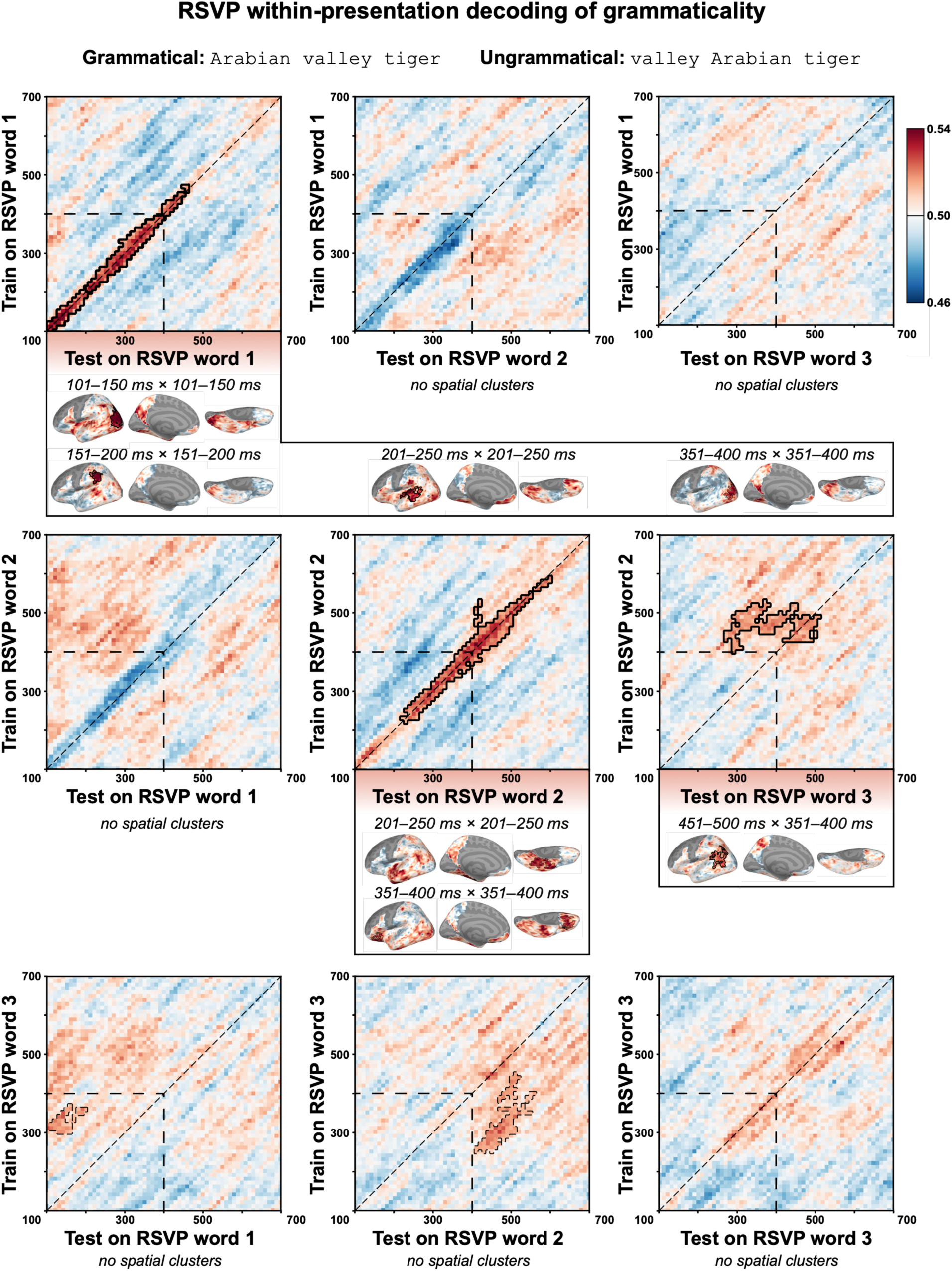
RSVP within-presentation decoding of grammaticality. Classifiers trained and tested on the same word position were successful at words 1 and 2, but not at word 3. Some generalization to other word positions was observed when trained on RSVP word 2 or RSVP word 3. Spatial decoding revealed recruitment of occipital cortex as well as more classical language areas. Temporal train-test time windows were 100–400 ms and 100–700 ms. Solid cluster outline: *p* < 0.05; dashed cluster outline: 0.05 < *p* < 0.1. RSVP: Rapid Serial Visual Presentation.

#### Within-presentation decoding: RPVP

In the RPVP condition, an early cluster emerged from the 100–400 ms window (train: ∼100–310 ms; test: ∼100–280 ms; *p* = 0.0142. Expanding the analysis window to the 100–700 ms window revealed an additional later cluster (train: ∼440–590 ms; test: ∼350– 650 ms; *p* = 0.0346). Both clusters had some off-diagonal distribution, indicating temporal generalization across train-test data. The early temporal cluster yielded a spatial cluster within the 251–300 ms × 101–150 ms tile (*p* = 0.0125), recruiting posterior-ventral parts of the temporal lobe, while the later temporal cluster returned an occipital spatial cluster within the 551–600 ms × 601–650 ms tile (*p* = 0.0489).

#### Within-presentation decoding: RSVP word 1 training

Training on RSVP word 1 and testing on RSVP word 1 produced a sustained, diagonal cluster in the 100–400 ms window (train: ∼100–400 ms; test: ∼100–400 ms; *p* = 0.007), which extended even further in the 100–700 ms window (train: ∼100–470 ms; test: ∼100– 460 ms; *p* = 0.0197). Several spatial clusters arose from this cluster, moving from occipital areas in the 101–150 ms × 101–150 ms tile (*p* = 0.0125) over the inferior parietal lobule in the 151–200 ms × 151–200 ms tile (*p* = 0.0444) to posterior-middle parts of the temporal lobe in the 201–250 ms × 201–250 ms tile (*p* = 0.0332) and, finally, back to occipital areas with a marginal cluster in the 351–400 ms × 351–400 ms tile (*p* = 0.0539). Temporal classifiers trained on RSVP word 1 and tested on either RSVP word 2 or word 3 did not produce significant temporal clusters (100–400 ms: all *p* ≥ 0.56; 100–700 ms: all *p* ≥ 0.56).

#### Within-presentation decoding: RSVP word 2 training

While no clusters resulted from RSVP word 2-training and RSVP word 1-testing (100– 400 ms: all *p* ≥ 0.68; 100–700 ms: all *p* ≥ 0.13), testing on RSVP words 2 and 3 both yielded significant temporal clusters. Similar to what was seen for RSVP word 1 train-test, RSVP word 2 train-test produced a sustained, diagonal cluster (train: ∼220–400 ms; test: ∼220–400 ms; *p* = 0.0106), with the 100–700 ms revealing its full extent (train: ∼220–590 ms; test: ∼220–600 ms; *p* = 0.0083). This temporal cluster resulted in two marginal spatial clusters: One with an anterior-ventral temporal localization in the 201–250 ms × 201–250 ms tile (*p* = 0.071) and another localizing to ventromedial prefrontal cortex in the 351–400 ms × 351–400 ms tile (*p* = 0.0743). Testing on RSVP word 3 yielded a late cluster with substantial temporal generalization in the 100–700 ms window (train: ∼400–560 ms; test times ∼250–510 ms; *p* = 0.0148), with the spatial decoding analyses returning a single cluster localizing to posterior-ventral temporal areas in the 451–500 ms × 351–400 ms tile (*p* = 0.0391).

#### Within-presentation decoding: RSVP word 3 training

Training on RSVP word 3 and testing on each of the three word positions only produced marginal clusters. Specifically, testing on RSVP word 1 produced a brief marginal cluster in the 100–400 ms window (train: ∼300–370 ms; test times ∼100–200 ms; *p* = 0.089) whereas testing on RSVP word 2 produced a longer and broader marginal cluster in the 100–700 ms time window (train: ∼250–450 ms; test times ∼410–570 ms; *p* = 0.0537). We did not perform spatial decoding analyses within these temporal clusters given that they did not reach significance. Unlike what was seen for within-word position testing for RSVP words 1 and 2, training and testing on RSVP word 3 did not produce any clusters (100– 400 ms: all *p* ≥ 0.19; 100–700 ms: all *p* ≥ 0.31)

## DISCUSSION

In this direct comparison of RSVP and RPVP, we found behavioral and neural evidence for shared encoding of phrase grammaticality (*Arabian valley tiger* vs. *valley Arabian tiger*) across presentation modes, although these similarities were also accompanied by some differences. In our behavioral matching task, grammatical phrases had higher accuracies and faster reaction times compared to ungrammatical phrases (though the latter remained a trend for RPVP), consistent with prior reports on the faciliatory effect of well-formed structure (Pegado et al., 2021; Fallon & Pylkkänen, 2024; Flower & Pylkkänen, 2024; Dunagan et al., 2025; Krogh & Pylkkänen, 2025; Li & Pylkkänen, 2026). Such sensitivity to grammaticality arose despite inherent differences in stimulus delivery timing and input structure, as evidenced by the main effects of presentation mode: On average, RPVP trials elicited faster reaction times while RSVP trials had higher accuracy. Notably, the use of novel noun-noun compounds presented in all caps suggests that grammaticality detection does not depend on conceptual familiarity or recognizable visual envelopes cuing different word categories. Thus, behavioral sensitivity to grammaticality appears to arise from linguistic computations rather than some form of superficial mapping to phrasal templates.

By training a classifier on neural data from one presentation mode and evaluating its performance on the other, we directly assessed the representational invariance of grammaticality. This produced some evidence for shared neural patterns across RPVP and RSVP, suggesting common underlying mechanisms that distinguish grammatical from ungrammatical phrases. Specifically, our temporal generalization analyses produced significant clusters at the second word in both train-test directions, albeit with different latencies and duration, alongside an early, off-diagonal cluster when training on RPVP and testing on RSVP word 3. That cross-presentation generalization is possible is remarkable given the vast differences between RPVP and RSVP, proving that the neural data are sufficiently similar to support cross-presentation generalization. Nevertheless, the temporal divergences for clusters obtained at RSVP word 2 and the unidirectional decodability at RSVP word 3 at the same time suggests that the two presentation modes appear to differentially encode different processes.

These presentation-invariant representations likely reflect (the output of) linguistic computations rather than low-level visual encoding for three reasons. First, the lexical material at the first word varies across grammatical and ungrammatical phrases in exactly the same way as at the second word (cf. the contrast between *Arabian* and *valley* at both word positions in *<u>Arabian</u> <u>valley</u> tiger* and *<u>valley</u> <u>Arabian</u> tiger*), yet effects were only obtained at the latter. Second, we observed an effect at the third word despite identical lexical material across conditions (*Arabian valley <u>tiger</u>* vs. *valley Arabian <u>tiger</u>*), providing evidence that this shared representation must reflect the composite of combinatory operations. Finally, follow-up spatial decoding analyses identified areas in the left anterior temporal lobe rather than in occipital areas. While our spatial decoding analyses are inherently limited in their ability to capture the distributed information relied upon by temporal decoding analyses—where individual sources may contribute to pattern discriminability without reaching significance in isolation—the identification of extra-occipital activity nevertheless supports a linguistic rather than perceptual explanation. This is particularly relevant for analyses of the second word as this word was presented foveally in the RPVP condition.

In fact, the involvement of specifically left anterior temporal areas when training on RSVP word 2 and testing on RPVP provides strong evidence that this effect reflects linguistic processes. Prior MEG-RSVP studies, many using a two-word minimal compositions paradigm, have consistently found elevated activation in the left anterior temporal lobe when words compose together, across visual and auditory modalities (e.g., Bemis & Pylkkänen, 2011, 2013, 2013) and irrespective of language and orthography switching (Phillips & Pylkkänen, 2021). Comparable patterns have likewise been observed during language production across speech and sign (Pylkkänen et al., 2014; Blanco-Elorrieta & Pylkkänen, 2017). Our localization of RSVP word 2-RPVP cross-presentation decoding to this particular region thus reinforces the hypothesis that the left anterior temporal lobe subserves core linguistic processes regardless of how language is externalized, even if our effect emerged later (at ∼500 ms) than is typically seen in RSVP research. Although several of our within-presentation spatial analyses also implicated the left temporal lobe, the cross-presentation activity was more focal, potentially because cross-presentation decoding isolates purely linguistic processes unconstrained by the temporal structure of the input. Why our spatial decoding analyses did not yield effects for the reverse train-test direction remains a question for future research; however, following the reverse logic provided for the lack of occipital clusters, this null result does not preclude potential left anterior temporal lobe involvement.

In our complementary within-presentation decoding analyses, we found that grammaticality was generally decodable when training and testing on data from the same presentation mode and, in the case of RSVP, word position. Follow-up spatial decoding analyses found that such effects recruited a wide range of brain areas, ranging from occipital cortex to areas widely recognized to support language processing such as large swaths of the temporal lobe and ventromedial prefrontal cortex. As such, linguistic processes rather than purely perceptual ones must have been at play. Nevertheless, the lack of within-presentation temporal decodability when training and testing on RSVP word 3— where the lexical material was identical across the grammatical and ungrammatical conditions—does suggest that differences in visual percept may aid cluster formation.

Interestingly, the nature of the within-presentation clusters was different across presentation modes: While training and testing on RPVP produced broad early and late clusters, training and testing on either of RSVP words 1 or 2 produced sustained clusters firmly anchored to the diagonal. Such differences could indicate that the higher load of the RPVP condition introduces greater inter- and intrasubject variability in processing than does RSVP, resulting in the greater extent of temporal generalization at the early and late clusters and the lack of decodability between 300–500 ms.

### Bidirectional cross-presentation grammaticality encoding at word 2 reflects different processes

The successful, bidirectional decoding at the second word suggests that the signal at this stage is dominated by presentation-invariant operations or representations, independent of the stimulus’ temporal structure. Although grammaticality is predictable by the first word, the second word represents the critical juncture where grammatical and ungrammatical phrases differ in their relative ease of composability. Consequently, the cross-presentation sensitivity to grammaticality likely reflects combinatorial processing of the first and second words.

Nevertheless, the different latencies and durations of clusters across the two train-test directions suggest that they reflect different aspects of presentation-invariant operations or representations. When comparing our temporal cross-presentation results to the within-presentation ones, the effect of training set becomes clear. For example, the cluster obtained when training on RSVP word 2 and testing on RPVP lasted from ∼300–500 ms, which falls squarely within the within-presentation RSVP word 2 cluster but outside the bounds of the within-presentation RPVP clusters. In contrast, the cluster arising from training on RPVP and testing on RSVP word 2 began much earlier and showed a temporal train-test shift, largely encompassed by the early and broad within-presentation RPVP cluster.

Delineating the nature of what the clusters reflect is more challenging, not least because our stimuli’s unusual grammaticality violation (correct vs. incorrect order of adjectival and nominal modifiers) and use of similarity-rather than difference-based methodologies complicate comparisons with prior literature. One possibility for the RSVP word 2-RPVP result is that it corresponds to N400 component observed in EEG-RSVP studies employing violation paradigms; for a review see Kutas & Federmeier (2011). This would align well with the localization to temporal regions, even if our result is more anterior than often observed for N400 components (Lau et al., 2008).

The lack of spatial localization makes interpretation of the RPVP-RSVP word 2 cluster even harder, but the rapidity of the grammaticality sensitivity does resemble what has been observed in some prior MEG-RPVP work (Fallon & Pylkkänen, 2024; Li & Pylkkänen, 2026). Intriguingly, Li & Pylkkänen (2026) obtained their early results from an adjective order contrast (*tiny green leaf* vs. *green tiny leaf*), thus being very similar to our modifier grammaticality manipulation. Building on evidence suggesting some degree of left-to-right processing even when a full stimulus is presented in parallel (Flower & Pylkkänen, 2024), Li & Pylkkänen speculate that the left edge of a stimulus presented with RPVP receives early processing priority, which would therefore lead to processing differences between stimuli where one modifier more readily initiates combinatory operations than the other. Since our grammaticality manipulation swapped the order of adjectival and nominal modifiers and because adjectives regularly serve an attributive function and therefore appear in combinatory contexts, our results could support such a hypothesis rooted in degree of combinatorial ease. Naturally, however, nouns can likewise serve as modifiers of other nouns, just like adjectives can be used on their own as predicates, thus emphasizing the need for future targeted investigations of a left edge processing priority.

### Unidirectional cross-presentation grammaticality encoding at word 3

Unlike what we saw at the second word, cross-presentation decoding at the third word was asymmetric, succeeding only when training on RPVP and testing on RSVP, but not vice versa. The resulting cluster had delayed test times (∼230–350 ms) relative to train times (∼130–250 ms), notably overlapping almost perfectly with the RPVP-RSVP word 2 cluster. This raises questions about why decoding may succeed in one direction but fail in the other. Since the third word marks the completion of stimulus presentation in RSVP, the shared representation may reflect the rapid encoding of global structure as also seen in prior MEG-RPVP work (Fallon & Pylkkänen, 2024). We hypothesize that such global structure is encoded more robustly in RPVP than in RSVP because the simultaneous presentation of all three words enables immediate structure building or, perhaps, detection (see, e.g., Fallon & Pylkkänen, 2024; Dunagan et al., 2025; Flower & Pylkkänen, 2026a). Conversely, the slow unfolding of stimuli in RSVP may cause processes partially or fully absent in the RPVP representation to dominate the signal and therefore disrupt generalization in the reverse direction. We speculate that phrasal “wrap-up effects”, reflecting delayed consolidation of the third word with the previous ones (Fodor & Bever, 1965; Just & Carpenter, 1980), could be one such process masking global structure encoding in the RSVP signal, but leave it to future research to explore this.

### Absence of cross-presentation grammaticality encoding at word 1

What might explain the lack of shared grammaticality encoding at the first word, despite it being the first cue to ungrammaticality? The successful RSVP word 1 within-presentation decoding clearly indicates that participants detected this lexical distinction:

Grammatical phrases began with a geographical or material modifier (e.g., *<u>Arabian</u> valley tiger* or *<u>plastic</u> valley tiger*) whereas ungrammatical phrases began with a location or animal modifier (e.g., *<u>valley</u> Arabian tiger* or *<u>tiger</u> Arabian valley*). The absence of cross-presentation decoding therefore suggests an absence of comparable word 1 encoding within the RPVP signals.

This could plausibly have two origins. First, consistent with the logic outlined regarding the unidirectional effects at the third word, the RPVP signal may be dominated by, for example, the global structure, thus overshadowing the retrieval of individual lexical items. Alternatively, or as an additive factor, the first word of RPVP stimuli may simply have been processed less deeply than the second and third words. In a left-to-right language like English, the perceptual span extends further to the right than to the left (for reviews, see Rayner, 1998, 2009), potentially disadvantaging the first word despite all stimuli falling within the range of parafoveal vision and our behavioral analysis of mismatch trials not indicating a disadvantage for word 1. Crucially, this interpretation does not contradict the potential for a left edge processing advantage as discussed in light of the word 2 cross-presentation results: Because the combinatory context only emerges upon the presentation of RSVP word 2, the shared representation of grammaticality required for cross-presentation decoding does not yet exist at RSVP word 1. At present, we cannot rule out either hypothesis.

## CONCLUSIONS

To conclude, we have demonstrated that sensitivity to grammaticality across Rapid Serial Visual Presentation (RSVP) and Rapid Parallel Visual Presentation (RPVP) shares core behavioral and neural characteristics, yet also diverges in subtle ways. While grammatical phrases consistently enjoyed a behavioral processing advantage over ungrammatical phrases, overall speed and accuracy differed significantly between the two presentation modes. Neurally, our decoding analyses revealed most pronounced evidence of similarity at the point where (un)grammaticality occurred, not when it could be predicted. In some cases, we found train-test directions to matter for decodability; for example, decoding was successful when training on RPVP and testing on RSVP word 3, but not vice versa. Such asymmetries suggest that while the neural data likely contain reflexes of the same underlying mechanisms, the two presentation modes may differentially prioritize certain aspects of the signal. That shared representations emerge at all is noteworthy since RPVP simultaneously integrates multiple lexical items whereas RSVP is rooted in incremental, serial processing. Because effects observed in one presentation mode were not always mirrored in the other, RSVP and RPVP are best viewed as complementary rather than interchangeable methodologies. If used in tandem, they may provide a more comprehensive characterization of the neural underpinnings of language processing than either paradigm alone.

## ACKNOWLEDGEMENTS

This work was supported by the National Science Foundation award #2335767 (LP). We thank Bernarda Basualdo for assistance with data collection as well as Alec Marantz, Ailís Cournane, Sebastian Michelmann, Alona Fyshe, and the members of NeLLab for their suggestions and feedback.

## COMPETING INTERESTS

The authors declare no competing interests.

## AUTHOR CONTRIBUTIONS

Conceptualization: SK, LP, Methodology: SK, LPm Investigation: SK, Data curation: SK, Formal analysis: SK, Software: SK, Resources: SK, Validation: SK, LP, Visualization: SK Funding acquisition: LP, Project administration: SK, LP, Supervision: LP, Writing – original draft: SK, Writing – review & editing: SK, LP

## SUPPLEMENTARY TABLE LEGENDS

**Table S1:**
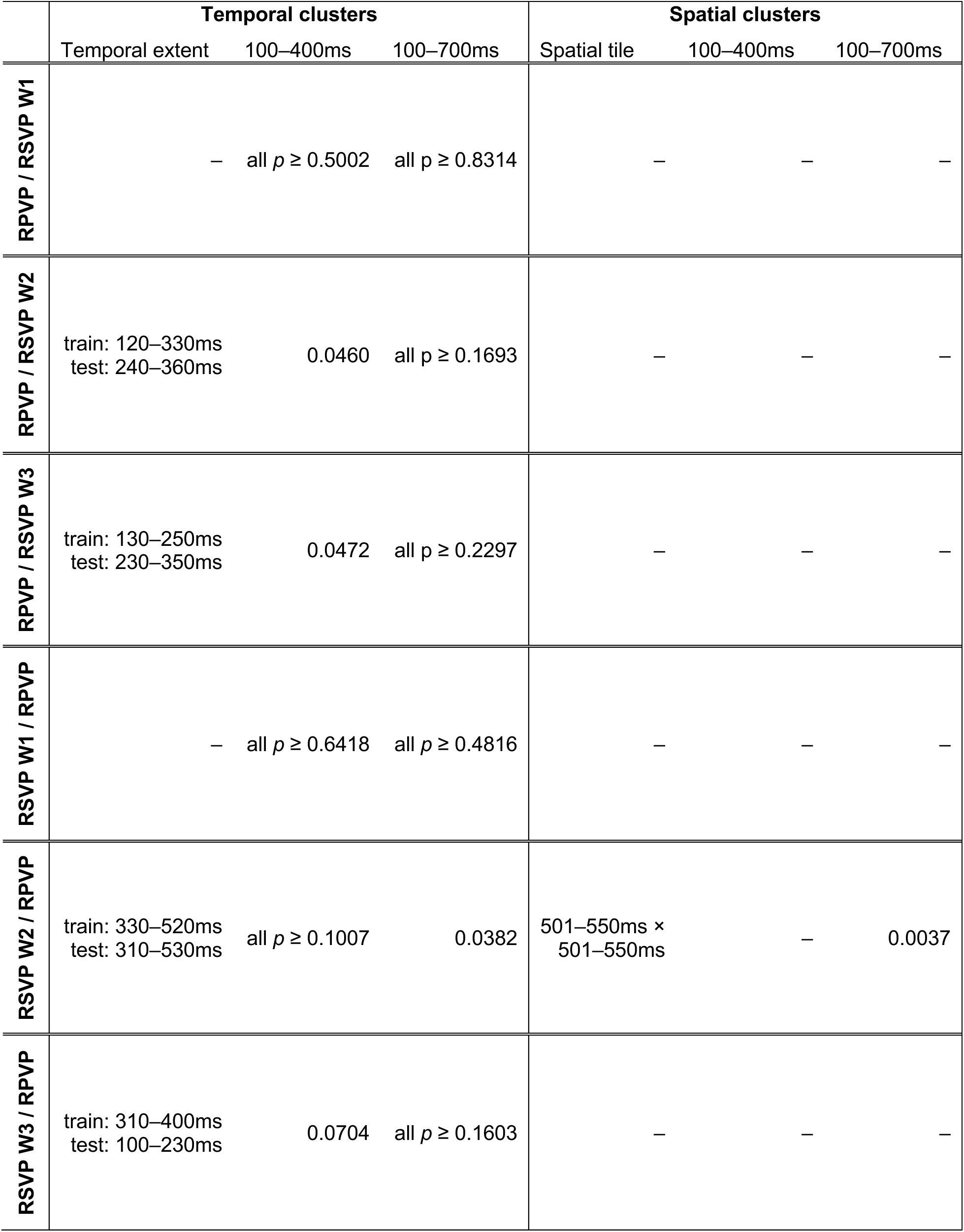
Cross-presentation temporal and spatial clusters. Significant clusters from the 100–400 ms and 100–700 ms windows with their corresponding *p* values and any spatial clusters. RPVP: Rapid Parallel Visual Presentation; RSVP: Rapid Serial Visual Presentation.

**Table S2:** RPVP within-presentation temporal and spatial clusters. Significant clusters from the 100–400 ms and 100–700 ms windows with their corresponding *p* values and any spatial clusters. RPVP: Rapid Parallel Visual Presentation.

|  | **Temporal clusters** |  |  | **Spatial clusters** |  |  |
| --- | --- | --- | --- | --- | --- | --- |
|  | Temporal extent | 100–400ms | 100–700ms | Spatial tile | 100–400ms | 100–700ms |
| **RPVP / RPVP** | train: 100–310ms<br>test: 100–280ms | 0.0142 | 0.0198 | 251–300ms ×<br>101–150ms | 0.0125 | 0.0125 |
|  | train: 440–590ms<br>test: 350–650ms | – | 0.0346 | 551–600ms ×<br>601–650ms | – | 0.0489 |

**Table S3:**
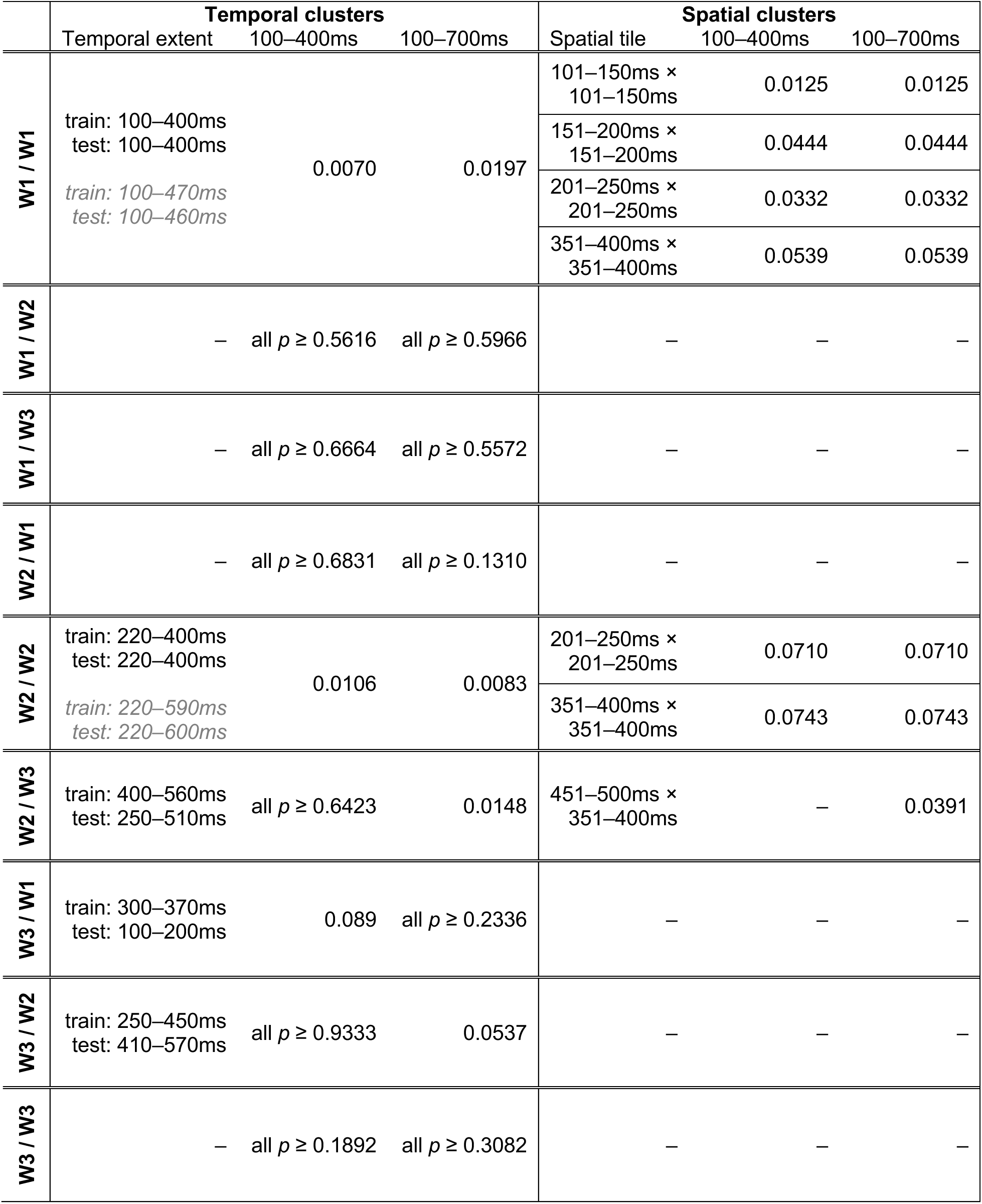
RSVP within-presentation temporal and spatial clusters. Significant clusters from the 100–400 ms and 100–700 ms windows with their corresponding *p* values and any spatial clusters. Grey text indicates full extent of clusters observed in the 100– 400 ms. Grey italicized text indicates extended cluster duration as revealed by the 100– 700 ms window. RSVP: Rapid Serial Visual Presentation.

## Notes

https://osf.io/hx85b/overview

### Competing Interest Statement

The authors have declared no competing interest.

